# Hoop-like role of the cytosolic interface helix in *Vibrio* PomA, an ion-conducting membrane protein, in the bacterial flagellar motor

**DOI:** 10.1101/2021.10.27.466211

**Authors:** Tatsuro Nishikino, Yugo Sagara, Hiroyuki Terashima, Michio Homma, Seiji Kojima

## Abstract

*Vibrio* has a polar flagellum driven by sodium ions for swimming. The force-generating stator unit consists of PomA and PomB. PomA contains four-transmembrane regions and a cytoplasmic domain of approximately 100 residues which interacts with the rotor protein, FliG, to be important for the force generation of rotation. The three-dimensional structure of the stator shows that the cytosolic interface (CI) helix of PomA is located parallel to the inner membrane. In this study, we investigated the function of CI helix and its role as stator. Systematic proline mutagenesis showed that residues K64, F66, and M67 were important for this function. The mutant stators did not assemble around the rotor. Moreover, the growth defect caused by PomB plug deletion was suppressed by these mutations. We speculate that the mutations affect the structure of the helices extending from TM3 and TM4 and reduce the structural stability of the stator complex. This study suggests that the helices parallel to the inner membrane play important roles in various processes, such as the hoop-like function in securing the stability of the stator complex and the ion conduction pathway, which may lead to the elucidation of the ion permeation and assembly mechanism of the stator.

**Importance:** Bacteria have a motor embedded in the membrane to rotate flagella as screw for swimming. The motor is composed rotor and stator complexes. The interaction between the rotor and stator converts the electrochemical potential gradient across the membrane into motor torque. The stator functions as an ion channel and is composed of two membrane proteins, MotA and MotB for proton or PomA and PomB for sodium ion. Based on the structural data of stator, we systematically introduce the proline replacement mutations and found that the cytosolic interface (CI) helix which is located parallel to the inner membrane between the second and third transmembrane (TM) segments, performs a hoop-like function in securing the stability of the stator complex and the ion conduction pathway. The results of this study provide novel insights into the energy conversion mechanism of the flagellar motor and the general mechanism of the ion channel function.

## Introduction

Most motile bacteria have filamentous macromolecular machine called flagella that extend outside the cell and rotate like a screw to move to a favorable environment for survival. An ion-driven rotary motor consists of a stator and a rotor at the base of the flagellum. The stator converts the electrochemical potential difference between the inside and outside the cell into a torque that is generated by interaction with the rotor. The coupling ions that drive the flagellar motor are known to be protons (H^+^) in *Escherichia coli* and *Salmonella*, or sodium ions (Na^+^) in *Vibrio* species and alkalophilic *Bacillus* (1-3).

The stator is composed of MotA and MotB in *E. coli* and *Salmonella*, and the orthologs PomA and PomB in *Vibrio*. PomA (MotA) is a four-transmembrane (TM) protein with a large cytoplasmic loop (Loop_2-3_) between the second and third TM regions. The loop_2-3_ contains several conserved charged residues, and the electrostatic interactions between these residues and the conserved charged residues of FliG of the rotor are important for motor rotation (4-7). PomB (MotB) is a single TM protein and the TM region is present in the N-terminus and the C-terminal region is present in the periplasmic space (8, 9) In the periplasmic region of PomB (MotB), there is an OmpA-like domain responsible for peptidoglycan binding (PGB), which anchors the stator to the peptidoglycan layer around the rotor via a PGB motif and the PGB region of Pal is interchangeable with the PGB region of MotB (10, 11). The crystal structures of the PGB region have already been solved (12-14), and it has been speculated that the structure changes significantly when the stator unit is activated, allowing it to be assembled and anchored around the rotor periphery (15). A segment of approximately 15 residues, called the plug region adjacent to the MotB or PomB TM region, prevents the ion influx of the stator as when the stator does not assemble around the rotor (16-18).

The structure of the cytoplasmic loop (Loop_2-3_) of the stator PomA (MotA) interacting with the rotor (FliG) is very important for the mechanism of torque generation (5, 6). The interaction between the PomA cytoplasmic loop and the FliG C-terminal region was directly detected by the site-directed photo-crosslinking between the residues of PomA D85, R88, K89, G90, F92, L93, or E96 and the residues of the FliG C-terminal region (19). We have previously attempted to clarify the structure of Loop_2-3_ by preparing various constructs. However, they all precipitated when overexpressed in *E. coli*, and we could not proceed to the structural analysis (20). Thus, we predicted the structure of PomA based on the structure of ExbB, whose structures have been reported (21, 22) and showed weak amino acid sequence similarity to PomA. Furthermore, ExbB is a membrane protein complex responsible for energy conversion using an ion-driven force and has an operating mechanism similar to that of PomA. Based on the ExbB atomic structure, we predicted the stator structure composed of PomA and PomB, and Pro-substituted mutants were constructed in Loop_2-3_. Last year, the structures of MotAB stator derived from *Campylobacter jejuni, Clostridium sporogenes*, and *Bacillus subtilis* were determined by single-particle analysis using cryo-electron microscopy (23-25). The MotA/MotB and PomA/PomB complexes, previously proposed as 4:2 hetero-hexamers, were shown to be 5:2 hetero-heptamers. From the structure in which two molecules of MotB are inserted in the center of the MotA ring made of five molecules, a model was proposed in which the MotA ring rotates with respect to the axis of MotB due to the influx of ions. It has been recently suggested that the plug region functions as a “spanner” to prevent the stator PomA pentamer ring rotation around the PomB TM axis, so that the ion flux and the stator rotation are coupled (26).

In this study, we focused on the characteristic cytosolic interface (CI) helix parallel to the inner membrane interface in Loop_2-3_. To investigate the roles of this helix in the stator function, we made Pro and other amino acid substitutions in the CI helix of PomA.

## Results

### Motility and protein expression of the Pro mutants

We predicted the secondary structure of PomA based on the structure of ExbB (22, 27), which showed homology to PomA, and introduced proline replacements, a predicted helix breaker, into the residues of the helix that follow the TM2 helix (G53 to A70) in the PomA Loop_2-3_ region (Fig. 1A). After construction, the stator structures were reported by single-particle analysis using cryo-electron microscopy (23, 25). Thus, we realized that the mutations were introduced in the CI helix of Loop_2-3_ parallel to the inner membrane (Fig. 1B and 1C). Plasmids expressing Pro-substituted PomA and wild-type (WT) PomB were introduced into NMB191 (*pomAB*-deficient strain) and expressed by arabinose induction. Motility was evaluated by swimming ring formation on a soft agar. The K60P, K64P, F66P, and M67P mutants showed no swimming ring formation, and the lack of motility was confirmed by light microscopy (Fig. 2A). The protein expression of each mutant was detected by western blotting using an anti-PomA antibody. All mutants, except for the K60P mutation, produced PomA (Fig. 2B). It is noteworthy that most mutant PomA bands were detected as a single band in western blotting, whereas the WT PomA was detected as a double band. This suggests that these mutations affect the structure of PomA.

**Fig. 1.**
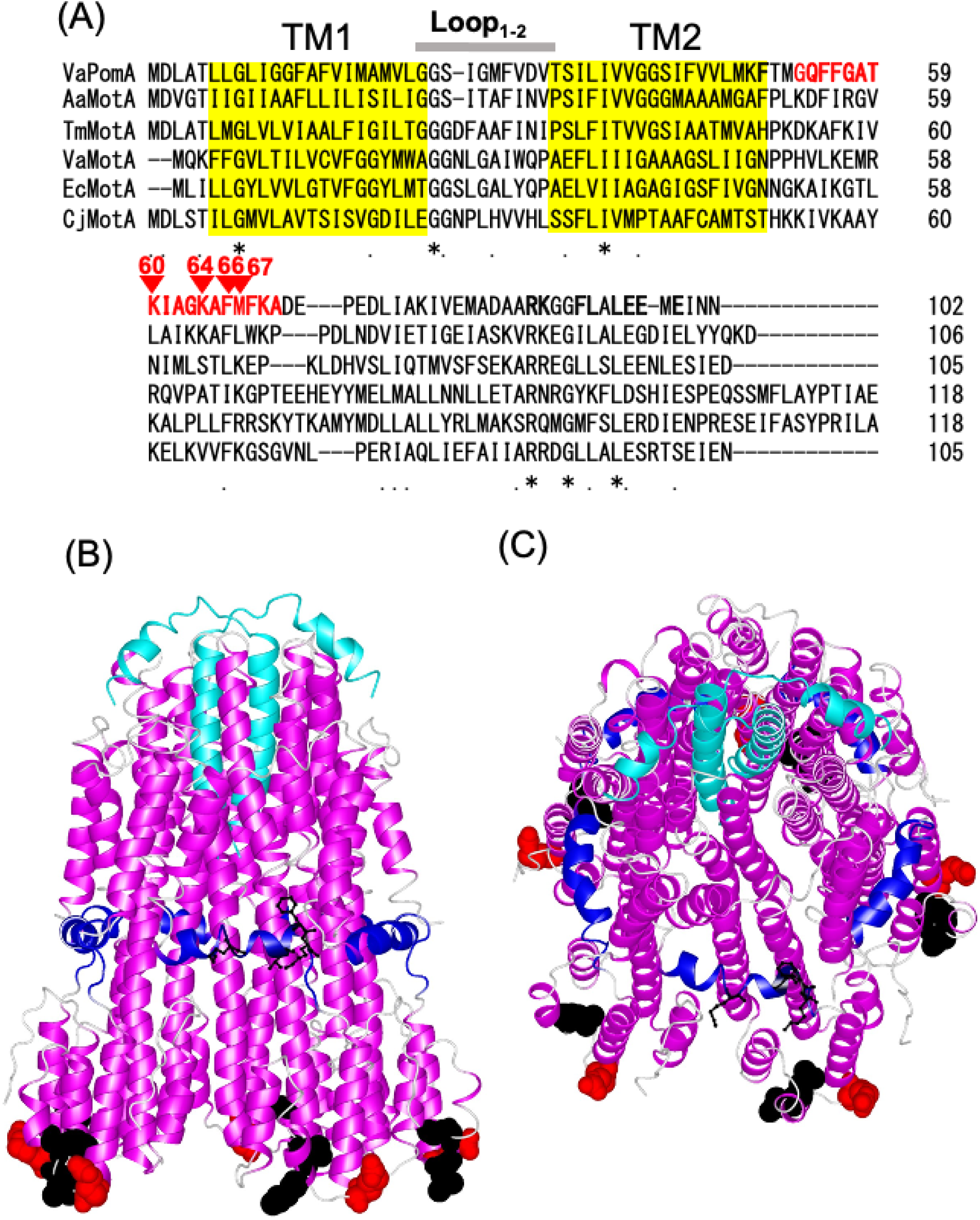
(A) Alignment of the N-terminal sequences of the stator proteins, PomA and MotA. The transmembrane region (TM) is shown in yellow. The mutated residues in the PomA of *Vibrio alginolyticus* mutated in this study are shown in red. VaPomA: PomA of *Vibrio alginolyticus* VIO5 (wild-type strain for the polar flagellum), AaMotA: MotA of *Aquifex aeolicus*, TmMotA: MotA of *Thermotoga maritima*, VaMotA: MotA of *Vibrio alginolyticus* VIO5, EcMotA: MotA of *Escherichia coli*, and CjMotA: MotA of *Campylobacter jejuni*. (B, C) Atomic structure model of the *C. jejuni* (Cj) stator, MotAB (PDB ID: 6YKM). The MotB part corresponding to PomB, and the cytosolic interface (CI) region of MotA corresponding to PomA are shown in light blue and blue, respectively. The residues of MotA corresponding to K60, K64, F66, and M67 of PomA are shown by the ball-and-stick model in black. The residues of MotA corresponding to the putative interacting charged and hydrophobic residues PomA are shown by the space filling model in red and black, respectively. (B) Side view of the stator and (C) slanting view from the top.

**Fig. 2.**
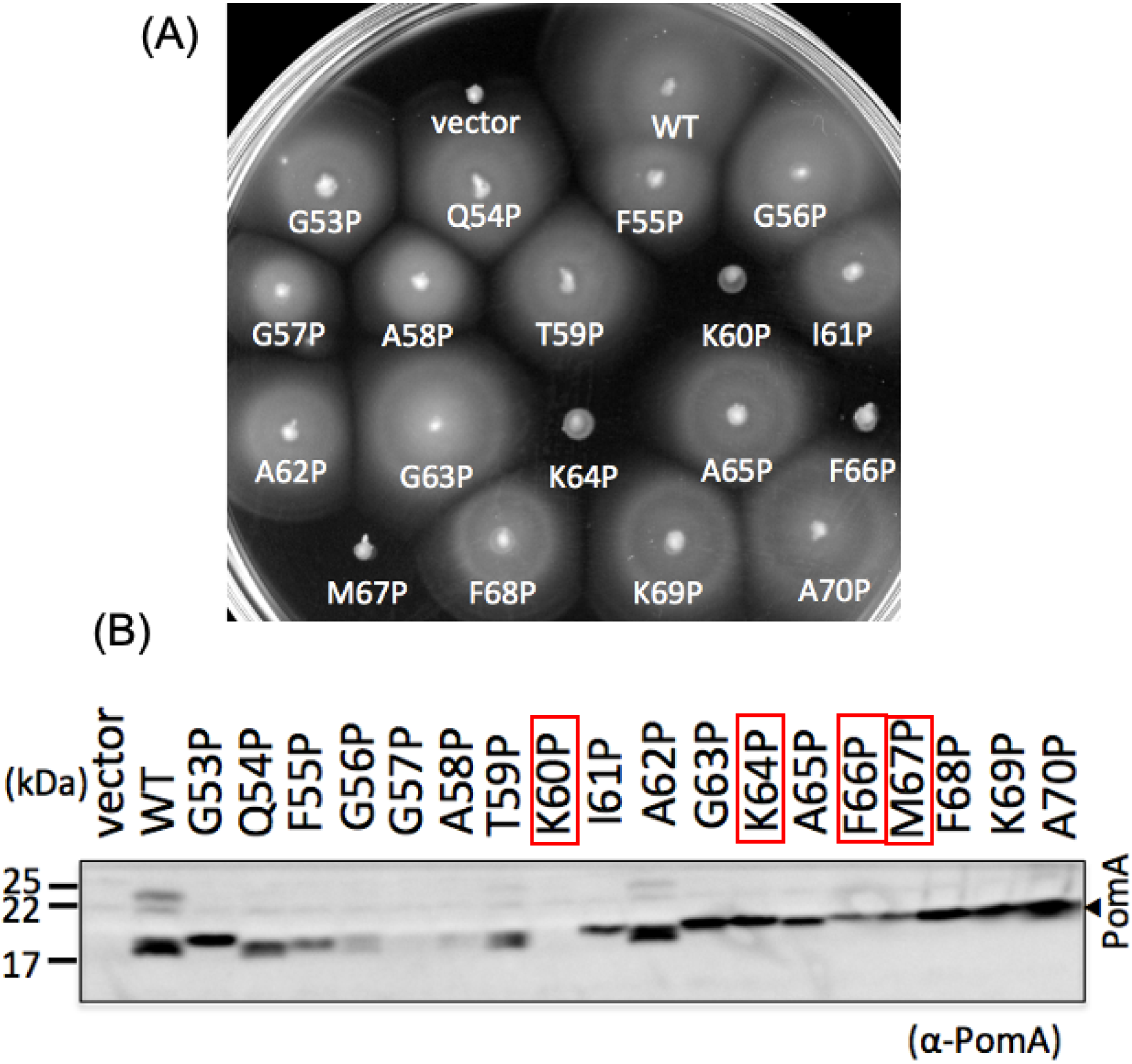
The motility of Pro substitution mutant and its protein expression. (A) *Vibrio* NMB191 (*pomAB* mutant) cells harboring the pHFAB-based plasmid with the *pomA* mutations from the fresh colonies were inoculated in soft agar plates (VPG 0.25%) with 0.02% arabinose and incubated at 30 ºC for 5 h. (B) *Vibrio pomAB* mutnat cells harboring the same pHFAB-based plasmid above with mutations were grown to the mid-log phase. The proteins of the cells were then separated using sodium dodecyl sulfate-polyacrylamide gel (SDS-PAGE) and detected via western blotting using the anti-PomA antibody.

### Motility of Ala and Ser mutants and dominant effects of the mutants

To investigate the residue specificity of the Pro substitutions, we replaced the residues K64, F66, and M67 with Ala or Ser. The motility assay on soft agar showed that the motility of K66A, M67A, and M67S was similar to that of the WT, and that of the K64A mutant was reduced, while that of the K64S and F66S mutants was completely lost (Fig. 3A). We speculate that the M67P mutation might affect the structure of the F66 residue, which is the neighbor of M67. Protein expression of each mutant was detected by western blotting using an anti-PomA antibody, and protein expression was confirmed in all mutants (Fig. 3C).

**Fig. 3.**
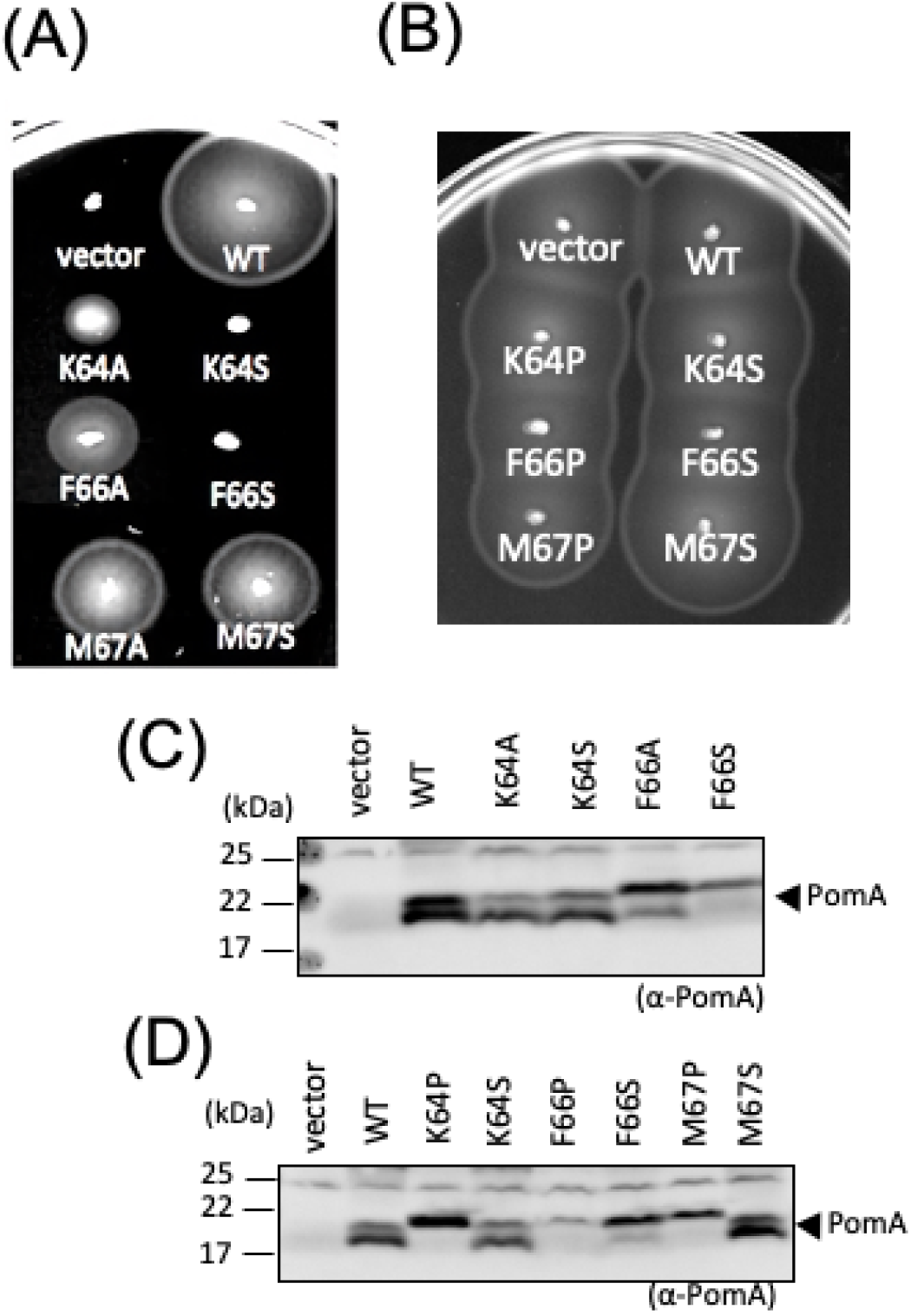
Mutations other than Pro and the dominant effects caused by the mutants. *Vibrio pomAB* (A) or wild-type (B) cells harboring the pHFAB-based plasmid without PomA (vector) or with wild-type PomA (WT) and the Ala and Ser substituted mutants (K64A, K64S, F66A, F66S, M67A, or M67S) from the fresh colonies were inoculated in soft agar plates (VPG 0.25%) with 0.02% arabinose and incubated at 30 °C for 5 h. *Vibrio pomAB* (C) or wild-type (D) cells harboring the same pHFAB-based plasmid above with the mutations were grown to the mid-log phase. The proteins of the cells were then separated using SDS-PAGE and detected by western blotting using an anti-PomA antibody.

Next, we examined the dominance of these mutants. A plasmid expressing the mutant PomA was introduced into the WT VIO5 strain of the polar flagellar motor, and PomA mutant expression was induced by arabinose. The plasmid-derived mutant PomA was expressed more than the chromosome-derived WT PomA. The dominant effects of Pro and Ser mutants were not observed (Fig. 3B and 3D).

### Analysis of stator function

The lack of the dominant effect of the mutant PomA suggests that the mutant stator complex is not able to assemble around the rotor. To examine whether Pro-substituted PomA does not assemble around the rotor, we expressed the mutant GFP-fused PomB with PomA from a plasmid in NMB191 (*pomAB*-deficient strain) by induction of arabinose. When WT PomA and GFP-fused PomB were expressed, fluorescence dots were observed at the poles of cells, indicating that the PomAB complex was assembled around the motor (Fig. 4A). On the other hand, almost no fluorescent dots could be observed in the PomA-K64P and F66S mutants, indicating that the PomAB complex cannot assemble around the rotor.

**Fig. 4.**
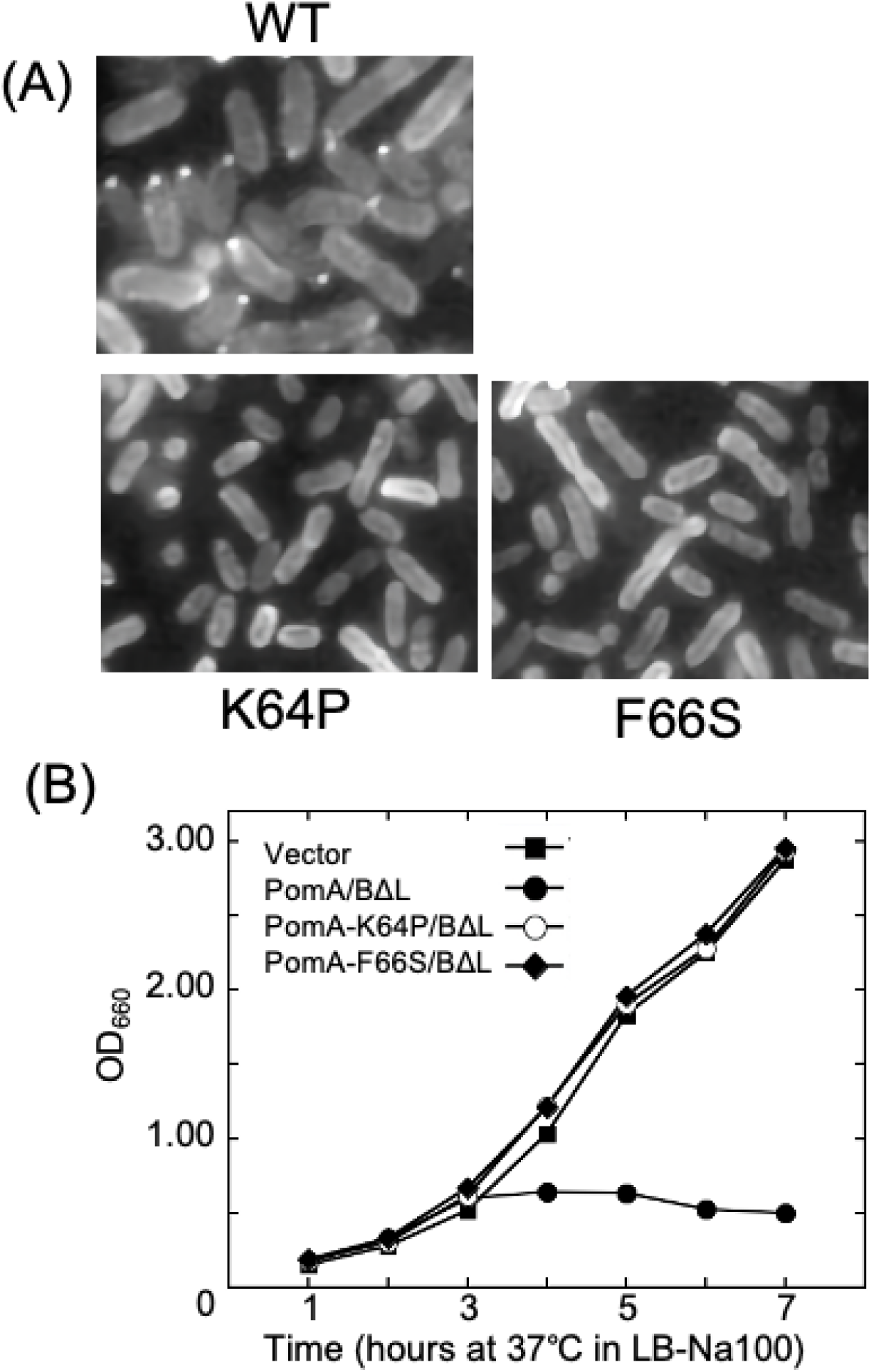
Profiles of the K64P and F66S mutants. (A) *Vibrio pomAB* cells harboring the pHFGBA with wild-type PomA (WT) or K64P and F66S of PomA substituted mutants were cultured in VPG broth containing 0.02% arabinose for 4 h at 30 °C and were observed by fluorescent microscopy. (B) Growth curve of cells. Overnight culture of *E. coli* cells harboring pBAD33 (Vector) and pTSK37 (PomA/BΔL) and derivative plasmids (PomA/BΔL D24N, PomA-64P/BΔL, and PomA-F66S/BΔL) was inoculated in the Luria–Bertani (LB) 3% sodium chloride (NaCl) broth at 1/100 dilution; arabinose was added at a final concentration of 0.02% (w/v), 2 h later, to induce expression (arrow). A_660_ was measured every 1 h after induction.

Although the loss of motility by the Pro mutation was explained by the inability of stator assembly around the rotor, we examined whether the mutant PomA with PomB, itself, is functional, that is, whether the mutant PomAB complex is capable of ion permeation. The plug region (residues 44-58) in PomB acts as a lid to prevent ion flow from the extracellular region. When the plug mutant (PomB_ΔL_) is expressed in *E. coli* cells, excessive ion influx by the stator inhibits cell growth. When mutant PomA and PomB deficient in the plug region were co-expressed from a plasmid in *E. coli* DH5α by arabinose induction, growth inhibition was suppressed by the K64P and F66S mutations (Fig. 4B). This suggests that the K64P and F66S mutations affect the structure of the ion channel.

### Purification of stator complex

We have improved the purification of the PomAB stator complex by using a cold-shock vector overexpression system in *E. coli* and the detergent, decyl maltose neopentyl glycol (DMNG), and have been able to obtain a reproducible, stable, and highly pure purified complex (28). We purified the PomAB stator with PomA F66S and K64P mutations. We could purify the mutant stator similar to the WT, although the amount of both mutants was reduced. After affinity chromatography and Coomassie brilliant blue (CBB) staining of the purified sample, bands of PomA and PomB were observed, confirming the formation of the complex. The affinity chromatography sample was then concentrated, and gel filtration chromatography was performed using a diluted sample of WT (Fig. 5A). F66S, and K64P stator were eluted at peak volumes of 11.19 mL and 11.33 mL, which are close to the WT peak volume of 11.26 mL. It is noteworthy that the F66S and K64P complexes eluted with a shoulder slightly earlier than the peak positions. We do not know the reason why the elution profiles are slightly different compared to the WT, although we speculate that the binding number of detergents is different. To confirm the stator structure of the purified stator, we observed the stator by electron microscopy (Fig. S1). The particles of the stator were observed, and we could not find any notable differences in particle shapes between the WT. Detailed structural analysis by cryo-electron microscopy is currently in progress.

**Fig. 5.**
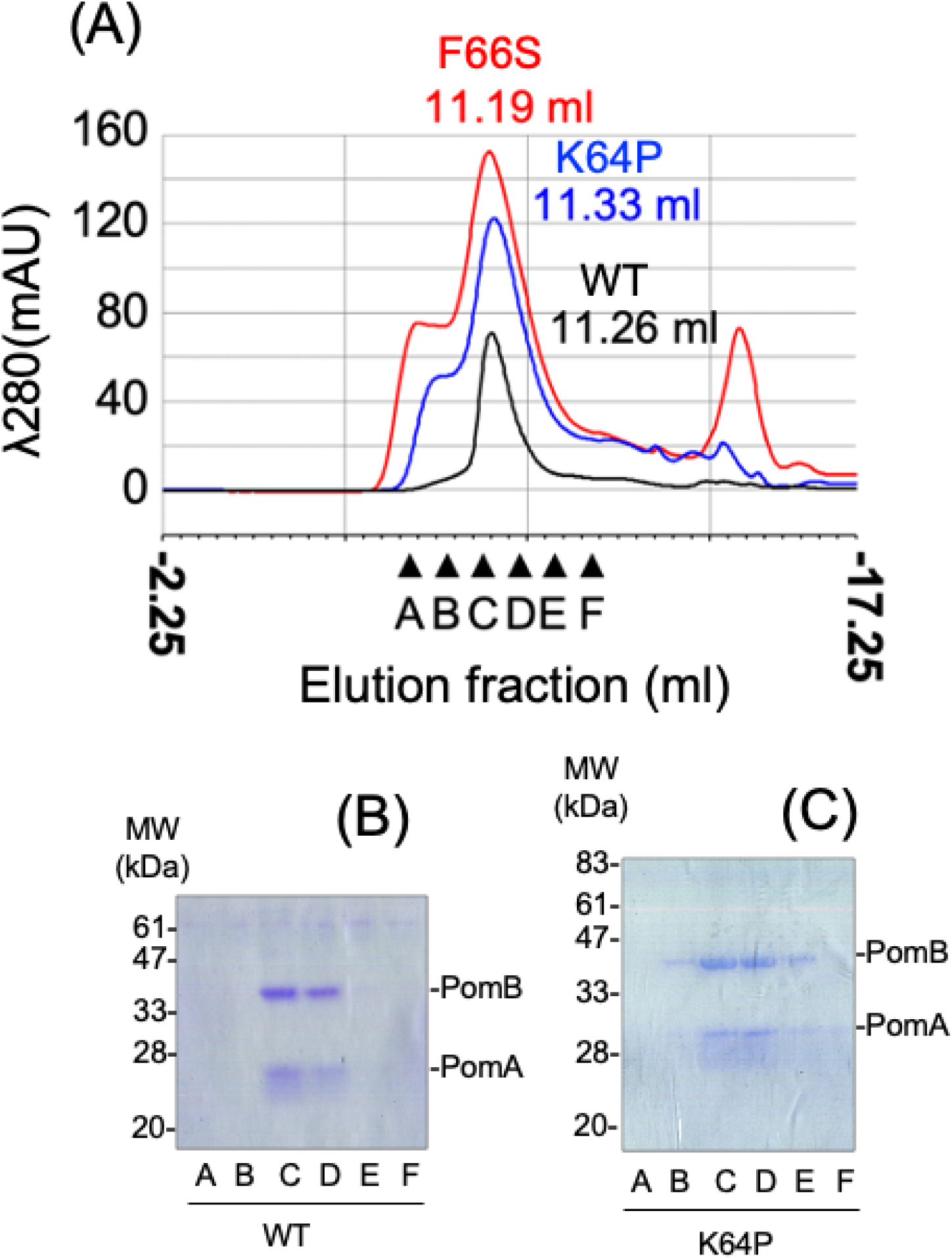
Profiles of the purified stator with the mutants in PomA. (A) Elution profile of the size exclusion chromatogram in the WT-stator complex and mutant. The stator expressed in *E. coli* cells by pColdIV-*pomA-pomB-his6* plasmid with WT PomA, K64P, or F66S mutation and purified by the affinity chromatography using hexa-histidine tag was subjected to size exclusion chromatography. The peak elution fractions (approximately at 10 mL) in each sample are shown. (B) Proteins of WT-stator, stator with K64P mutation in PomA, were purified and the elution samples by size exclusion chromatography were analyzed by SDS-PAGE and stained with Coomassie brilliant blue (CBB). The peak elution fractions (approximately at 10 mL) in each sample are shown.

### The cross-link formation between PomA TM1 and TM2

The CI helix lies between TM1 and TM2 and beneath the inner membrane and is located outside the pentamer as if each of it encircles a PomA molecule to hold the whole pentamer ring. Thus, we speculated that the CI helix might stabilize the molecules of PomA in the pentamer, and mutations in the CI region may destabilize the PomA structure, thereby losing the stator function. We investigated the residue pair of adjacent PomA molecules in the pentamer ring that are located at different distances to form a disulfide bridge when the residues are substituted with Cys (Fig. 6). We found that PomA with L22C and T33C mutations, which are close to each other in the pentamer ring, form multimers and appear to be up to pentamer, as judged by western blotting (Fig. 6A). Since the PomA-L22C/T33C mutant showed only slightly reduced motility compared to the WT or mutants with a single mutation in the soft agar plate (Fig. 6B), the stator function was not significantly affected by the disulfide cross-link. We expected that if the disulfide cross-link could stabilize the TM helices in the stator, then the function of the K66P or F66S mutant stator would be restored. We introduced the K64P and F66S mutations in the L22C and T33C mutant PomA. The disulfide cross-linking did not suppress the K64P and F66S mutations (Fig. 6D). The K64P and F66S mutations did not affect disulphide cross-link formation, suggesting that the structures of TM1 and TM2 were not affected by the K64P and F66S mutations (Fig. 6C).

**Fig. 6.**
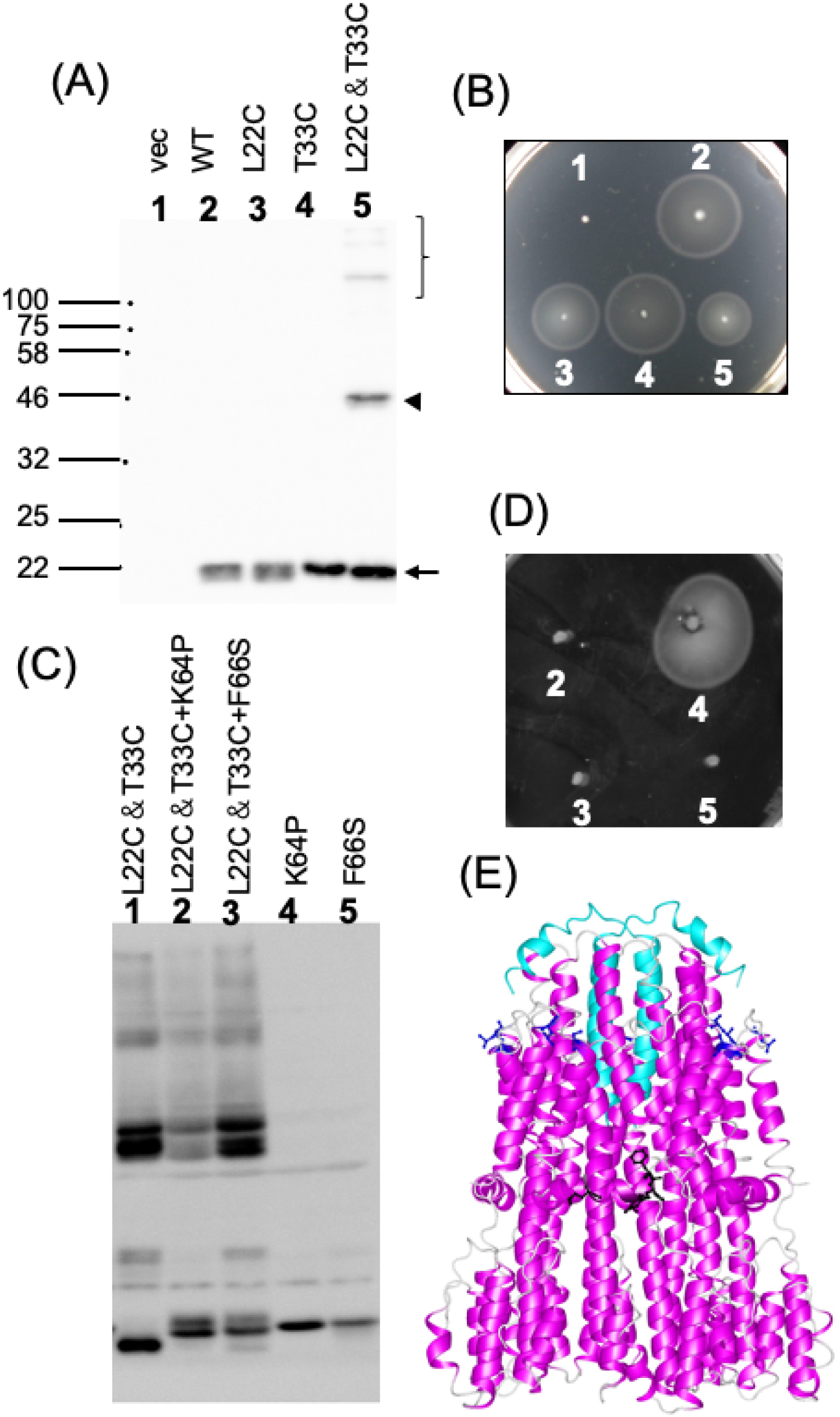
Effect of the PomA mutations on cysteine-substituted mutants. (A) Proteins extracted from *Vibrio pomAB* cells harboring 1: pBAD33 (vec), 2: pHFAB (WT), and pHFAB-based plasmid with the mutations, 3: *pomA-L22C* (L22C), 4: *pomA-T33C* (T33C), and 5: *pomA-L22C, T33C* (L22C&T33C) were separated using SDS-PAGE in the absence of a reducing agent. PomA was detected via western blotting using the anti-PomA antibody. (B) *Vibrio pomAB* cells harboring the same plasmids as (A) were inoculated in soft agar plates (VPG 0.25%) with 0.02% arabinose and incubated at 30 ºC for 5 h. (C) Proteins extracted from *Vibrio pomAB* cells harboring pHFAB-based plasmid with the mutations, 1: *pomA-L22C, T33C* (L22C&T33C), 2: *pomA-L22C, T33C, K64P* (L22C&T33C+K64P), 3: *pomA-L22C, T33C, F66S* (L22C&T33C+F66S), 4: *pomA-K64P* (K64P), and 5: *pomA-F66S* (F66S) were separated using SDS-PAGE in the absence of a reducing agent. PomA was detected via western blotting using the anti-PomA antibody. (C) *Vibrio pomAB* cells harboring the same plasmids as (D) were inoculated in soft agar plates (VPG 0.25%) with 0.02% arabinose and incubated at 30 ºC for 24 h.(E) Structure of *C. jejuni* (Cj) stator is shown by the ribbon model and the corresponding residues of *Vibrio* L22C and T33C mutations were shown in blue with the ball-and-stick model.

## Discussion

The flagellum is the locomotor machinery of many bacteria, including marine species belonging to the genus *Vibrio*. The rotational force is generated by the interaction between the stator and rotor via the ion flow in the stator. The interaction between the cytoplasmic region of Loop_2-3_ in the A subunit and the C-terminal region of the rotor protein of FliG is important for torque generation. We attempted to obtain the structural information of the Loop_2-3_ region, which is unknown. Last year, the three-dimensional structure of the stator complex (MotAB) from *Campylobacter jejuni, Clostridium sporogenes*, and *Bacillus subtilis* had been clarified (23, 25). We studied the sodium-driven flagellar motor of marine *Vibrio, Vibrio alginolyticus*. In this study, we systematically constructed proline-substituted mutants (from G53 to A70) in the cytoplasmic region close to TM2 in PomA. When the structure of MotA, which had been determined, was assigned to the corresponding structure and sequence of PomA, the Pro mutations were located in the CI helix, which is parallel to the inner membrane boundary (Fig. 1).

Among the 28 proline mutants, only three mutants (K64P, F66P, and M67P) were found to lose motility, even though the Pro mutant PomA proteins were expressed. These three residues (K64, F66, and M67) are localized in the C-terminal site of the CI helix, which is parallel to the inner membrane interface. To examine the effects of other amino acid substitutions, we introduced Ala or Ser into the residues K64, F66, and M67 (Fig. 3). We found that in the K64 residue, the Ala and Ser mutations reduced and lost motility, respectively, suggesting that the electrostatic interaction of these residues might be required for torque generation. In the F66 residue, the Ala and Ser mutations resulted in normal motility and loss of motility (Fig. 3 and S2), respectively.

Hydrophobicity or α-helix formation by this residue may be important for stator function. Other mutations did not affect motility. The WT PomA band by sodium dodecyl sulfate-polyacrylamide gel (SDS-PAGE) was detected as double-banded, suggesting that distinct stable structures were present even in the solution containing the strong detergent SDS. Most proline mutants of PomA in the CI helix become single-banded. This region may contribute to the stability of monomer PomA; however, the character of the monomer stability does not seem to correlate with the stator functions.

According to the dominant effect and the ability of poler localization of the stator complex, it seems that the CI mutant stators, which lost their function, could not assemble around the rotor (Fig. 4A). The stator assembly around the motor is necessary for activation and interaction with the peptidoglycan (PG) layer or T-ring depending on Na^+^ ions or interaction with FliG (29-32). The mutations may have altered the structure of the stator complex to prevent the sensing of Na^+^, thus inhibiting the interaction with the rotor. Next, we tested whether the stator could permeate ions in the growth assay using PomB lacking the plug region. PomA mutations suppressed growth inhibition (Fig. 4B), indicating that these mutations prevent ion flux. In other words, the structural instability caused by the mutations may abolish the rotor-stator interaction and block the Na^+^-conductive activity induced by the stator activation due to the interaction with the rotor. We have not yet clarified whether the mutation actually inhibits the ion permeation pathway or what kind of structural changes are induced by the mutations. Although the ion permeation pathway has not been clarified, the third and fourth TM regions of PomA and the TM region of PomB are thought to form the ion pathway, the PomB-D24 residue in the TM region and PomA-T158 and PomA-T186, which are located in TM3 and TM4, respectively, are known to be the essential Na^+^ binding sites (28, 33). Structural alteration at the C-terminal side of the CI helix may affect the structure of the Na^+^-binding site composed of these residues, although the overall structure of the stator complex was not altered by the mutations, as judged by gel-filtration chromatography of purified mutant stators (Fig. 5A, 5 B, and Fig. S1). It should be noted that the amount of expression was lower than that of the WT, and the shape of the gel filtration chromatography peaks did not allow us to conclude that this was due to structural differences.

The CI helix parallel to the inner membrane interface contains a large number of hydrophobic residues, suggesting that the hydrophobic profile of this helix serves as a CI helix. The F66 residue is highly conserved among the residues of the CI helix (Fig. 1A) and is likely to play an important role. Since the F66 residue is a hydrophobic residue with an inward-facing side chain, it may stabilize the structure through hydrophobic interactions with the other helices. The K64 residue is predicted to be exposed on the surface of the stator complex (Fig. 7 and S3). The K64 residue is a charged residue, and the electrostatic map of the stator complex (Fig. 7A) shows that the boundary region to the inner membrane, where the K64 residue is located, is positively charged, and the membrane is positively charged. This suggests that the K64 residue is involved in the stability of the stator through electrostatic interaction with the inner membrane, or is involved in the stabilization of the structure by interacting with charged residues in the helix extending from TM3 and TM4. In accordance with this assumption, the K203 residue, which is located at the C-terminal end of TM4 and structurally close to the K64 residue, is known to lose its motility due to the K203E mutation (34), suggesting that charged residues near the CI helix are important for stator function. The amino acid sequences of the CI region are not highly homologous among the species, although many negatively charged amino acids are present in the CI region and the CI helix are similarly arranged in the stator structures (Fig. S4). Based on our results, we propose that the CI helix, which lies parallel to the inner membrane, has a hoop-like function to support the TM3 and TM4 helices, which extend to the ion binding site. Interaction with the membrane surface with CI helices stabilizes the stator complex and ion-conducting pathway.

**Fig. 7.**
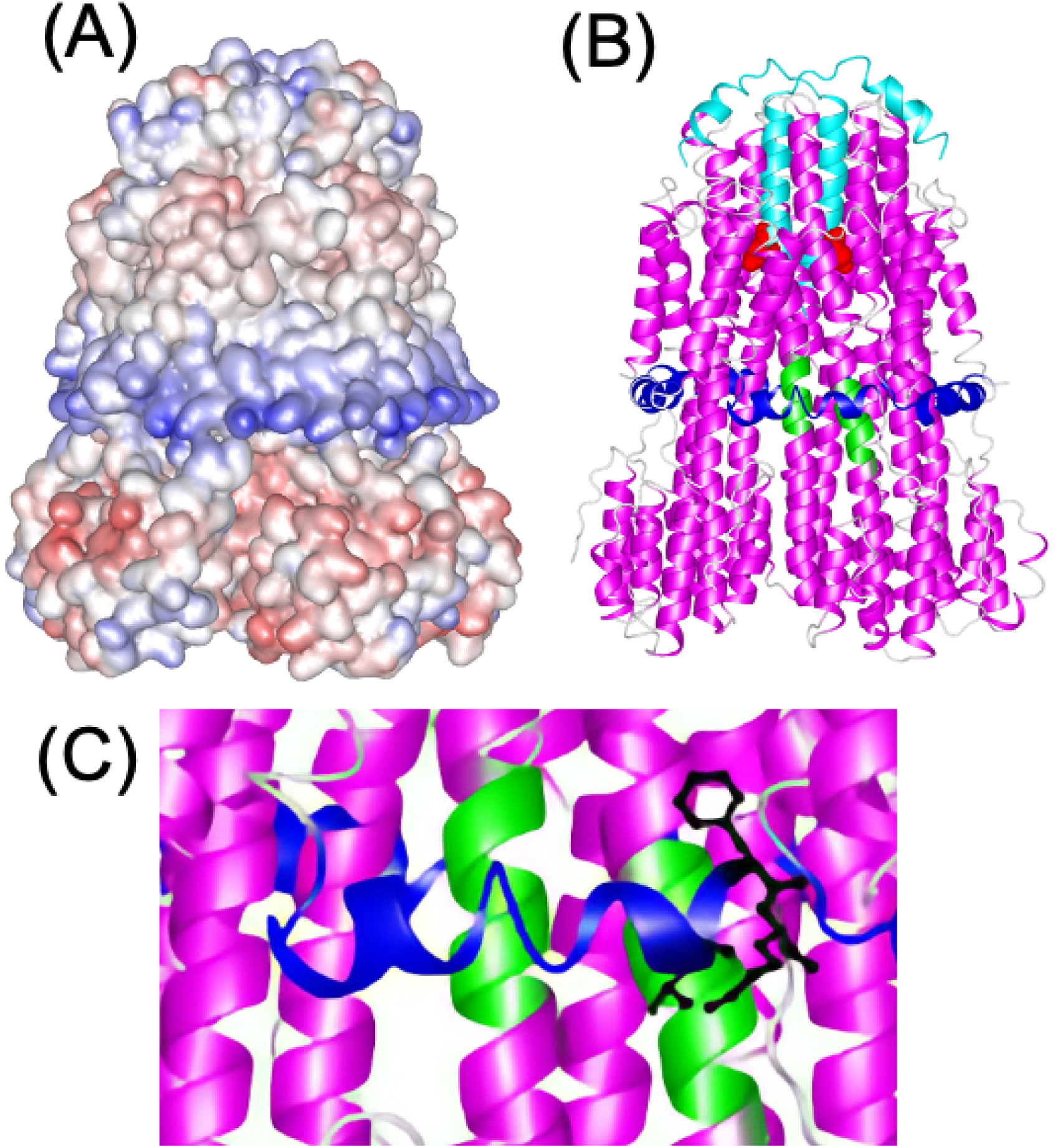
Possible function of the CI region in loop_2-3_ of *Vibrio* PomA. (A) Electrostatic potential map was estimated. (B) The structure of the Cj stator was shown by the ribbon model. The regions corresponding to G53-A70 of PomA and the B subunit are shown by blue and light blue, respectively. (C) The corresponding region of G53-A70 in (B) is expanded.

## Materials and Methods

### Bacterial strains and plasmids

*E. coli* was cultured in Luria–Bertani (LB) broth [1% (w/v) bactotryptone, 0.5% (w/v) yeast extract, and 0.5% (w/v) sodium chloride (NaCl)], LB 3% NaCl broth [1% (w/v) bactotryptone, 0.5% (w/v) yeast extract, and 3% (w/v) NaCl], and TG broth [1% (w/v) bactotryptone, 0.5% (w/v) NaCl, and 0.5% (w/v) glycerol). Chloramphenicol was added at a final concentration of 25 µg/mL for *E. coli*. Ampicillin was added at a final concentration of 100 µg/mL for *E. coli. V. alginolyticus* was cultured at 30 °C in VC medium [0.5% (w/v) polypeptone, 0.5% (w/v) yeast extract, 0.4% (w/v) K_2_HPO_4_, 3% (w/v) NaCl, and 0.2% (w/v) glucose) or VPG medium [1% (w/v) polypeptone, 0.4% (w/v) K_2_HPO_4_, 3% (w/v) NaCl, and 0.5% (w/v) glycerol]. If needed, chloramphenicol was added at a final concentration of 2.5 μg/mL for *V. alginolyticus* culture.

### Disulfide crosslinking experiment

Cells harboring the pHFAB plasmids were cultured in the VPG medium containing arabinose at a final concentration of 0.02% (w/v) at 30 °C for 4 h with 100-fold dilution from the overnight culture. To examine disulfide crosslinking, the cells were collected by centrifugation and then suspended in SDS-loading buffer without β-mercaptoethanol. SDS-PAGE and immunoblotting were performed as previously described (26).

### Swimming assay in soft agar plates

*Vibrio* NMB191 cells harboring the pHFAB-based plasmids with the mutants or wild-type *pomA* and *pomB* were plated on VPG plates with antibiotics. A colony of *Vibrio* cells was inoculated onto VPG agar [0.25% (w/v) bactoagar] plates containing 0.02% (w/v) arabinose and incubated at 30 °C.

### Purification of PomAB complex

The purification protocols were modified as previously described (28). BL21 (DE3) cells carrying the plasmid pCold4 *pomApomB*-*his*_6_ and pLysS were grown overnight in 30 mL of LB medium at 37 °C, inoculated in 1.5 L of LB medium, and incubated at 37 °C. When the cell density at OD_660nm_ approximately reached 0.4, cells were incubated in iced water for 30 min. Isopropyl β-D-1-thiogalactopyranoside (IPTG) was added to a final concentration of 0.5 mM to induce overexpression of PomA and PomB-His_6_ proteins and cultured for 1 d at 15 °C. Cells were harvested by centrifugation and the weight of the cells was measured. Seven times the volume of Na-Pi buffer (50 mM Na-Pi pH 8.0, and 200 mM NaCl) was added and the cells were suspended. To break the cells, the suspension was placed in a French press (OHTAKE) at 1,000 kg/cm^2^. Unbroken cells were removed by low-speed centrifugation. The samples were ultra-centrifuged at 118,000 *× g* for 1 h. The same volume of Na-Pi buffer before the French press was added to the resultant pellet and stored at –30 °C for later use. The frozen samples (10 mL and 40 mL volume) in the WT and mutants, respectively, were thawed in a water bath and stirred. To solubilize the resultant pellet, 10% (w/v) DMNG was added to a final concentration of 0.5% (w/v) and stirred for 60 min at 30 °C. The insoluble material was removed by ultra-centrifugation (118,100 *× g* for 30 min). The resultant supernatant was mixed with 4 mg of Talon Metal Affinity Resin (Takara) equilibrated with wash buffer [50 mM Na-Pi pH 8.0, 200 mM NaCl, 10 mM imidazole, and 0.02% (w/v) DMNG) and incubated at 4 °C for at least 1 h in a polypropylene column by batch method. After eluting the supernatant in the column, 4 mL (1 column volume) of wash buffer was added to the column three times to wash the column. To elute the His-tag stator from the resin, two column volumes of elution buffer [50 mM Na-Pi (pH 8.0), 200 mM NaCl, 200 mM imidazole, and 0.02 % (w/v) DMNG] was added and eluted. The his-tag affinity purified stator was concentrated to 1 ml using a 100 K MWCO Amicon device (Millipore). The samples were subjected to size exclusion chromatography using Enrich SEC650 column (Bio rad) in SEC buffer [20 mM Tris HCl (pH8.0), 100 mM KCl and 0.0025% (w/v) 2,2-didecylpropane-1,3-bis-β-D-maltopyranoside (LMNG)]. We set the flow at 0.75 mL per min and fraction volume of elution at 0.5 mL. The peak fractions were collected, and the concentrations were measured by absorptiometry (ε= 50310) of A280 using a Nanodrop (Thermo Scientific).

### Sample preparation and data correction of negative staining images

Elution fractions of WT and PomA-F66S mutations were diluted in SEC buffer. Final concentration of the samples in the WT and F66S mutation were 3.4 and 5.3 ng/mL, respectively. A 5 μL solution was applied to a glow-discharged continuous carbon grid. The excess solution was removed using filter paper, and the sample was subsequently stained on the carbon grid with 2% ammonium molybdate. Electron microscopy images were recorded with an H-7650 transmission electron microscope (Hitachi) operated at 80 kV and equipped with a FastScan-F114 CCD camera (TVIPS, Gauting, Germany) at a nominal magnification of 80,000 ×.

## Acknowledgements

This work was supported in part by the Japan Society for the Promotion of Science (JSPS) KAKENHI [Grant Numbers 18K07108 and 21K07022 (to H.T.), A20J00329 (to T.N.), and 20H03220 (to M.H.)].

## Supporting information

Supplementary information associated with this article can be found online on the publisher’s website.

